# P2X7 receptor signaling is not required but can enhance Th17 differentiation

**DOI:** 10.1101/2021.03.19.435855

**Authors:** Yin Yang, Meaghan E. Killeen, Tina L. Sumpter, Alicia R. Mathers

**Author notes:** <u>**Correspondence:**</u> email. Address: W1156 Biomedical Sciences Tower, 200 Lothrop St, Pittsburgh, PA 15261. (A.R.M.).

## Abstract

The purinergic receptor P2X7 (P2X7R) is important in inflammasome activation and generally considered to favor proinflammatory Th17 immune responses. However, several studies utilizing P2rx7^-/-^ mice did not observe impaired inflammation, on the contrary, these reports demonstrate that P2rx7^-/-^ mice can have more sever inflammation compared to WT controls. To begin to clarify this discrepancy, the impact of P2X7R signaling on primary Th17 and Th1 cell responses was examined. Initially, utilizing a global knockout approach, we found that P2rx7^-/-^ mice develop comparable Th17 and Th1 responses to those of WT mice. However, indepth *in vitro* and *in vivo* investigations revealed differences in the immune response outcome depending on which cell type expresses P2X7R. In this regard, DC-specific P2X7R deficient chimeras demonstrate a comparable Th17 differentiation pattern with that of WT chimeras but display a significantly enhanced Th1 response. However, P2rx7^-/-^ T cells have a suppressed Th17 differentiation profile while developing a similarly enhanced Th1 response as observed in DC-specific P2X7R deficient chimeras. Finally, P2X7R expression induces more T cell death *in vivo*, attributed to its cytotoxicity, which results in a similar total number of WT Th17 and P2rx7^-/-^ Th17 cells and remarkably higher amounts of P2rx7^-/-^ Th1 cells. Collectively, our findings establish that P2X7R expression on CD4^+^ T cells is necessary for Th17 differentiation while inhibiting Th1 development.

## Introduction

Cytokines are critical for intercellular communication in the immune system during its development and active immune responses. In addition to its role as energy currency and intracellular second messenger, adenosine triphosphate (ATP), has recently been recognized as an intercellular mediator that orchestrates immune responses. In this regard, ATP, an alarmin released from stressed, wounded, or necrotic cells, is a damage-associated molecular pattern (DAMP) that acts as a danger-signal initiating inflammation, including innate and adaptive immunity. Among the ATP receptors, P2X7R is physiologically necessary for its functions in triggering inflammation and its wide expression in mouse and human tissues. Mechanistically, when P2X7R is gated with a low ATP concentration it allows Ca^2+^ and Na^+^ influx and K^+^ efflux, which is beneficial for cell growth and cellular activation. However, a high concentration of ATP and prolonged stimulation will induce P2X7R to form a large nonselective membrane pore, which is linked to cell death and is dependent on the C-terminus of P2X7R (1,2). In mice, the cell death-inducing function can also be triggered by NAD^+^ through ADP-ribosylation of P2X7R by ADP-ribosyltransferase 2 (ART2) on plasma membrane (3,4).

A prototypical function of P2X7R signaling is the activation of the NLRP3 inflammasome in macrophages and monocytes, which leads to Caspase-1 activation and the processing and secretion of mature IL-1β (1). IL-1β is a proinflammatory cytokine involved in both innate and adaptive immune responses and inflammatory diseases (5). In line with this, P2X7R has been implicated in many inflammatory diseases, including multiple sclerosis, colitis, rheumatoid arthritis, psoriasis, and glomerulonephritis (6), diseases which Th17 cells have been shown to play a dominant role in the immunopathogenesis (7). Furthermore, studies suggest that P2X7R can regulate the function and development of many T cell subsets, including Th1, Th17, Treg, and Tfh cells (1,4). Likewise, Borges da Silva et al. demonstrate that P2X7R also promotes the maintenance of memory CD8^+^ T cells (8). However, reports regarding P2X7R’s effect on Th17 differentiation are controversial. In this regard, studies provided evidence that P2X7R contributes to Th17 responses in murine colitis (9), murine arthritis (10), murine and human psoriatic skin inflammation (11,12), and human adipose tissue inflammation (13). Whereas, reports also demonstrate that P2rx7^-/-^ mice develop more severe inflammation in the intestine during *Citrobacter rodentium* (Th17 and Th1) or *Toxoplasma gondii* (Th1) infection, and in the central nervous system in a model of experimental autoimmune encephalomyelitis (EAE; Th17) (14–16). Thus, the specific role and function of P2X7R in Th17 responses remains unclear.

In an attempt to begin to clarify P2X7R’s impact on T cell differentiation, we utilized a genetic technique to separately examine the respective contribution of P2X7R expression on dendritic cells (DCs) and CD4^+^ T cells to the differentiation of Th1 and Th17 cells using classical *in vitro* and *in vivo* adaptive immune models. For a more precise measurement of the Th17 response, in *in vivo* studies, we adoptively transferred naïve OT-II T cell receptor transgenic CD4^+^ T cells (OT-II cells) to congenic mice, the mice were then immunized with OVA_323_-_339_ peptide in complete Freund’s adjuvant (CFA), and Th17 differentiation of OT-II cells were tracked. We found *in vitro* that P2X7R expression on DCs can promote Th17 development when exogenous BzATP, an ATP analog and P2X7R agonist, is provided, whereas in the context of an *in vivo* setting classical DC (cDC)-specific ablation of P2X7R did not impair Th17 development. Isolated P2X7R expression on T cells demonstrated that P2X7R is required for optimal differentiation of Th17 cells *in vitro*. However, *in vivo* P2X7R expression on T cells promoted cell death and therefore resulted in a comparable total number of Th17 cells in both WT and P2rx7^-/-^ mice. Similar to findings achieved under cDC-specific P2X7R deficiency, P2X7R knockout on T cells also enhanced Th1 development.

## Results

### Global knockout of P2X7R does not compromise antigen-specific Th17 responses

To evaluate the contribution of P2X7R to Th17 immune response, we compared the IL-17A response in WT mice and P2rx7^-/-^ mice to keyhole limpet hemocyanin (KLH), which induces substantial Th17 responses (17,18). Thus, mice were immunized with KLH/CFA for 7 days and then IL-17A and IFNγ production were measured in antigen-specific recall responses. The results demonstrate that P2rx7^-/-^ mice are capable of mounting an antigenspecific IL-17 response comparable to WT mice (Fig. 1A). There was also no difference in IFNγ production between the two mouse strains. To determine if this response is generalized to other antigens, a similar experiment was conducted using OVA_323-339_ peptide to immunize mice (Fig. 1B). Consistently, P2rx7^-/-^ mice and WT mice produced comparable amounts of IL-17A and IFNγ. To further scrutinize the ability of P2X7R to induce Th17 differentiation and to begin to address the controversial findings, we assessed the role of P2X7R expression separately on DCs and CD4^+^ T cells.

**Figure 1.**
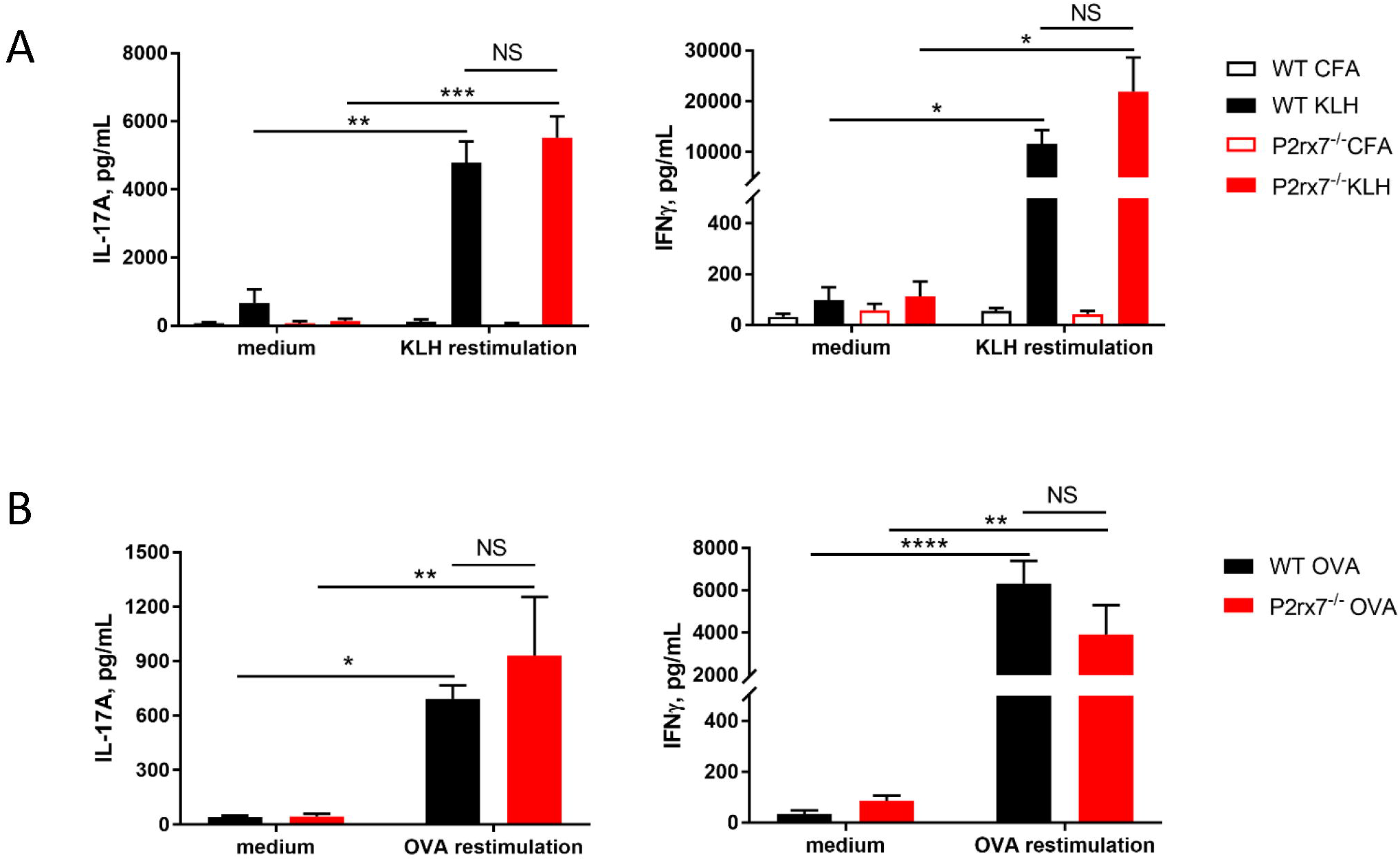
P2rx7^-/-^ mice mount comparable Th17 responses to WT mice. **(A)** WT C57BL/6J mice and P2rx7^-/-^ mice were immunized with 20 μg of keyhole limpet hemocyanin (KLH) emulsified in complete Freund’s adjuvants (CFA) for 7 days (KLH). Control mice were treated with CFA only (CFA). Inguinal lymph node (LN) cells were restimulated with KLH for 3 days and supernatants were collected for analysis of IL-17A and IFNγ by ELISA. **(B)** C57BL/6J mice and P2rx7^-/-^ mice were immunized with 20 μg OVA_323-339_ peptide emulsified in CFA for 7 days. OVA-specific IL-17A and IFNγ responses were analyzed as in **(A)**. Results shown as mean ± SEM. One representative of two independent experiments, n=6 total. *p<0.05, **p<0.01, ***p<0.001, ****p<0.0001; NS, not significant.

### *In vitro* stimulation of P2X7R on DCs enhances IL-17A production by allogeneic T cells

Our previous work has shown that human skin migratory DCs (mDCs) respond to P2X7R stimulation by producing proinflammatory cytokines and inducing the secretion of IL-17A from allogeneic T cells (11). To determine if mouse DCs have a similar response, we stimulated bone marrow-derived DCs (BMDCs) with varying doses of BzATP, an ATP analog and specific P2X7R agonist, and collected cell culture supernatants at the indicated time-points to examine the production of IL-6 and IL-1β, which are important for Th17 differentiation. In accordance with our previous data, we demonstrated that BzATP can significantly induce both IL-6 and IL-1β production by mouse BMDCs in a dose dependent manner (Fig. 2A). In addition, a mixed leukocyte reaction (MLR) was performed to see if mouse BMDCs stimulated with BzATP had the capacity to induce allogeneic Th17 differentiation. The results indicated that DCs pretreatment with BzATP significantly increased the production of IL-17A, compared to BMDCs not pretreated with BzATP (Fig. 2B). To demonstrate that BzATP-enhanced IL-17A production is dependent on DC expression of P2X7R, we expanded BMDCs derived from WT and P2rx7^-/-^ mice and performed a similar MLR. BzATP treatment of WT BMDCs significantly increased the IL-17A production, whereas BzATP had no effect on P2rx7^-/-^ BMDCs (Fig.2C). In the absence of exogenous BzATP, there was no significant difference between WT BMDCs and P2rx7^-/-^ BMDCs in IL-17A production, indicating that endogenous ATP is not contributing to the Th17 response (Fig. 2C). Collectively, these data indicate that BzATP stimulation of DCs can enhance IL-17A production of allogeneic T cells in a P2X7R dependent manner.

**Figure 2.**
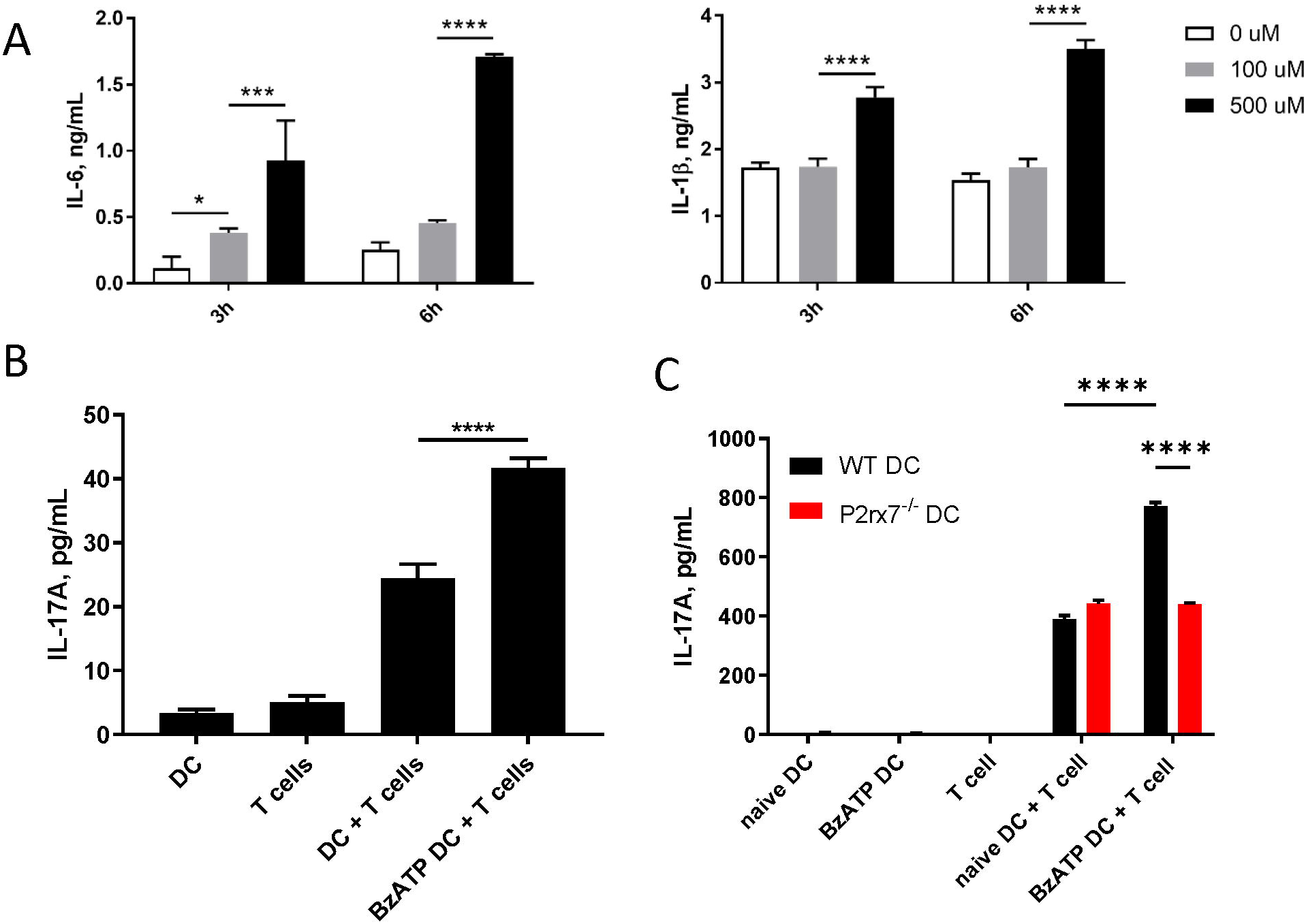
Stimulation of P2X7R on DCs can promote IL-17 production *in vitro*. **(A)** Mouse BMDCs derived from C57BL/6J mice were stimulated with the indicated doses of BzATP and supernatants were collected at 3h and 6h to determine IL-6 and IL-1β production by ELISA. **(B)** Mixed leucocyte reaction (MLR) was induced by cocultering BMDCs derived from C57BL/6J, pretreated with 300μM BzATP or left untreated (controls), with naive CD4^+^ T cells isolated from BALB/cJ for 4 days. Supernatants were collected and L-17A levels assessed by ELISA. **(C)** BMDCs derived from WT or P2rx7^-/-^ mice, pretreated with BzATP or left untreated (controls), were cocultured with naive CD4^+^ T cells from BALB/cJ for 4 days. Supernatants were collected to determine IL-17A by ELISA. Results shown as mean ± SEM. Results are one representative of three independent experiments. *p<0.05, ***p<0.001, ****p<0.0001; NS, not significant.

### *In vivo* P2X7R expression on classical DCs is dispensable for Th17 development

To assess the requirement of DC-expressed P2X7R for Th17 differentiation *in vivo*, we created an *in vivo* system in which P2X7R deficiency is restricted to only classical DCs (cDCs), utilizing the zDC-DTR mice (19). Zbtb46 (zDC) is a zinc finger transcription factor that is confined to the cDC lineage. Thus, diphtheria toxin (DT) treatment specifically ablates both CD8^+^ cDCs (CD103^+^ cDCs in nonlymphoid tissues) and CD11b^+^ cDCs subsets, while sparing monocyte-derived DCs, plasmacytoid DCs, macrophages, Langerhans cells, and NK cells. Utilizing the zDC-DTR mice we generated two groups of mixed bone marrow chimeras (Fig. 3A). Specifically, we reconstituted irradiated wild type C57BL/6J mice with bone marrow cells from WT mice and zDC-DTR mice (WT + zDC-DTR; 1: 1 ratio) or from P2rx7^-/-^ mice and zDC-DTR mice (P2rx7^-/-^ + zDC-DTR; 1:1 ratio). Following successful reconstitution chimeras were treated with DT to deplete cDCs (Fig. S1); thus, the cDC populations remaining are P2X7R deficient (cDC-specific P2X7R deficiency). We then utilized these chimeras to evaluate OVA-specific T cell development (Fig. 3A). Following re-stimulation, DT-treated P2rx7^-/-^ + zDC-DTR chimeras produced a comparable percentage of IL-17A^+^ OT-II cells to those recovered from DT-treated WT + zDC-DTR chimeras (Fig. 3, B and C). The absolute number of IL-17A^+^ OT-II cells in WT + zDC-DTR and P2rx7^-/-^ + zDC-DTR chimeras is also comparable; whereas, P2rx7^-/-^ + zDC-DTR chimeras induced a significantly higher percentage and absolute number of IFN*γ*^+^ OT-II cells P2rx7^-/-^ mice compared to WT mice. Taken together, these *in vivo* results indicate that P2X7R on cDCs is not required for Th17 differentiation but P2X7R signaling does block Th1 cell development.

**Figure 3.**
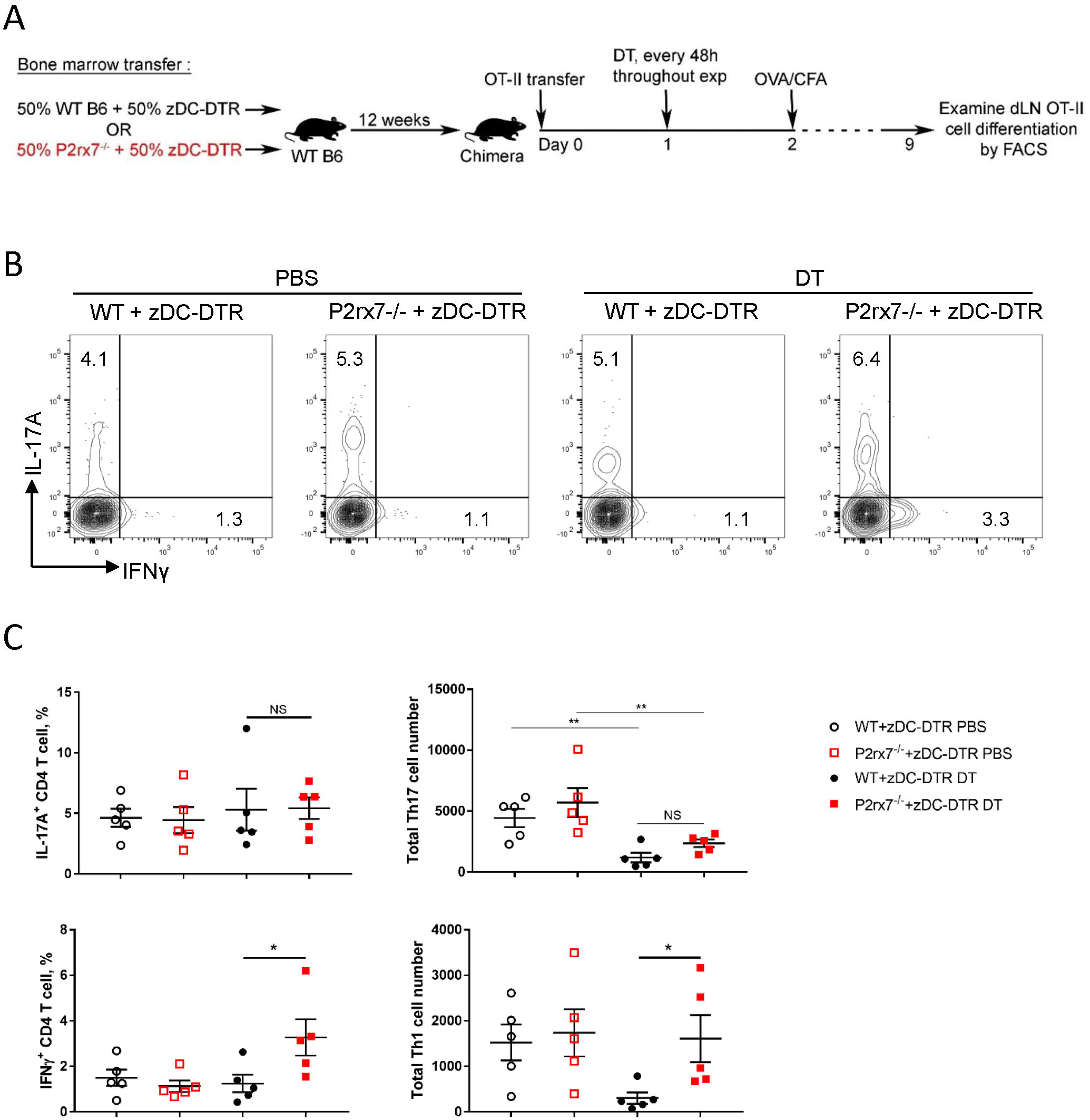
P2X7R on cDCs is not required *in vivo* for Th17 differentiation. **(A-C)** Mixed bone marrow chimeras were established by reconstituting irradiated WT mice with bone marrow from either WT or P2rx7^-/-^ mice and zDC-DTR mice mixed at 1:1 ratio (WT + zDC-DTR and P2rx7^-/-^ + zDC-DTR, respectively). After 10 weeks, chimeras were intravenously transferred with 1 × 10^5^ naive OT-II CD4^+^ T cells one day prior to injection with diphtheria toxin (DT) or vehicle. DT was injected every 48 h throughout the course of the experiment. 24 h following initial DT treatment, chimeras were immunized with OVA_323-339_/CFA, according to schematic **(A)** On day 9 inguinal LNs were collected, restimulated, and analyzed by flow cytometry. for the percentage of. **(B)** Representative flow contour plots. **(C)** The percentage of IL-17A^+^ and IFNγ^+^ OT-II T cells and cell number were quantitated. Results shown as mean ± SEM of two independent experiments, n=10 total. *p<0.05, **p<0.01; NS, not significant.

### P2X7R on T cells is necessary for Th17 development *in vitro*

It is possible that the P2X7R on T cells itself directly participates in Th17 differentiation patterns. Indeed, it has been reported that P2X7R is involved in T cell activation through regulation of the ATP autocrine pathway (20). To test this hypothesis, we adopted a Th17 differentiation coculture system (21). We co-cultured naive WT or P2rx7^-/-^ CD4^+^ T cells with WT or P2rx7^-/-^ BMDCs in the presence of anti-CD3ε, TGFβ, and LPS. P2rx7^-/-^ T cells demonstrated significantly impaired Th17 development but enhanced Th1 responses in terms of both the positive cell percentage and total cell number of IL-17^+^ and IFNγ^+^ cells (Fig. 4A and B). To confirm this conclusion, we next performed a cytokine-skewed Th17 differentiation assay to test the requirement of P2X7R on CD4^+^ T cells. We purified naive CD4^+^ T cells from either WT or P2rx7^-/-^ mice and stimulated them with anti-CD3 plus anti-CD28 in the presence of Th17-skewing cytokines IL-6, TGFβ, IL-1β, and IL-23. The P2rx7^-/-^ T cell treatment group demonstrated a significant decrease in the IL-17A^+^ cell percentage compared to WT T cells (Fig. 4C). Furthermore, the proliferation of P2rx7^-/-^ and WT T cells was comparable (Fig. 4D). Thus, together these *in vitro* results indicate that P2X7R expression on T cells is necessary for Th17 differentiation while inhibiting Th1 development.

**Figure 4.**
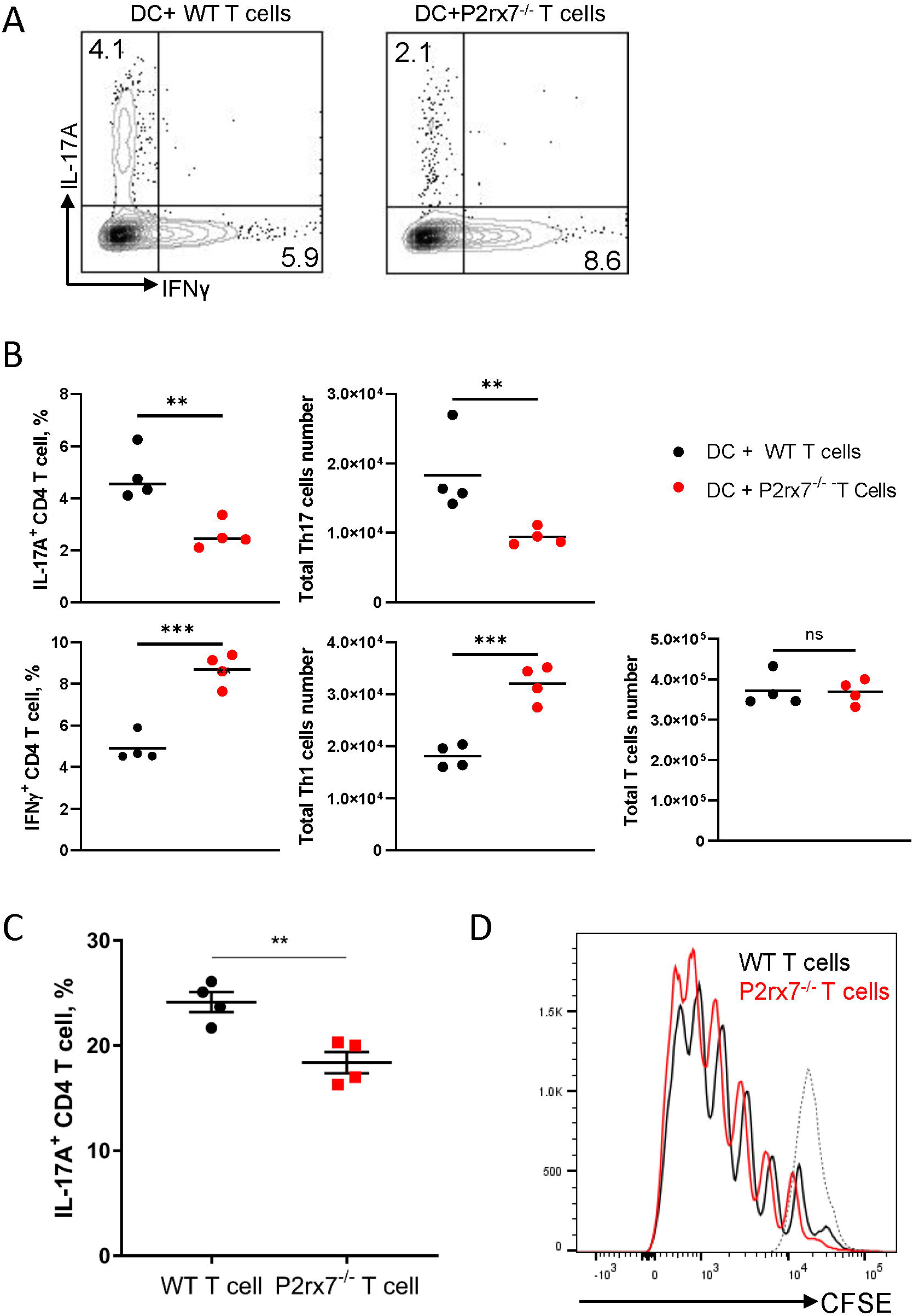
Stimulation of P2X7R on CD4^+^ T cells enhances Th17 development *in vitro*. BMDCs and naive CD4^+^ T cells from either WT or P2rx7^-/-^ mice were cocultured in the presence of 0.2 μg/ml anti-CD3, 100 ng/ml LPS, and 1 ng/ml TGF-β1 for 4 days. Cells were harvested, restimulated, and stained. IL-17A^+^ and IFNγ ^+^ CD4^+^T cell percentages were analyzed by flow cytometry. **(A)** Representative flow cytometric contour plots. **(B)** Percentage and total cell number of IL-17A^+^ and IFNγ^+^ CD4^+^T cells were quantitated. **(C)** Naive CD4^+^ T cells isolated from WT or P2rx7^-/-^ mice were stimulated with plate-bound anti-CD3 (10 μg/ml) and anti-CD28 (2 μg/ml) in the presence of Th17-skewing cytokines for 4 days. Percentages of IL-17A^+^ T cells were determined by flow cytometry following restimulation. **(D)** Naive CD4^+^ T cells isolated from WT or P2rx7^-/-^ mice, prelabelled with CFSE, were stimulated as in (C) for 4 days. T cell proliferation was analyzed by CFSE intensity in WT and P2rx7^-/-^ T cells. Dotted line represents CFSE labelled but unstimulated naive T cell. Results shown as mean ± SEM. A and B are one representative of five independent experiments. C and D are one representative of three independent experiments. *p<0.05, **p<0.01, ***p<0.001; NS, not significant.

### *In vivo* Th17 differentiation is not increased by P2X7R expression on T cells

To address the *in vivo* physiological relevance of P2X7R expression on T cells to the differentiation of Th17 cells, we generated P2rx7^-/-^ OT-II mice and examined adoptively transferred WT and P2rx7^-/-^ OT-II T cell differentiation in congenic wild type mice. As demonstrated in Figure 5A and B, P2rx7^-/-^ OT-II cells differentiated to IL-17A^+^ cells at a significantly lower percentage compared to WT OT-II cells, whereas the percentage of IFNγ^+^ cells was significantly higher in the P2rx7^-/-^ OT-II population. Of note though, the total number of P2rx7^-/-^ OT-II cells was higher than that of WT OT-II cells (Fig 5B), which resulted in equivalent total number of IL-17A^+^ cells in P2rx7^-/-^ and WT OT-II populations and an increase in the number of IFNγ^+^ cells. Because P2X7R has been shown to induce T cell death (14,22,23), we utilized a viability dye to assess the cell death of OT-II cells (Fig. 5C). Consistent with these previous reports, WT OT-II cells demonstrated a much higher percentage of cell death compared to P2rx7^-/-^ OT-II cells, which lead to a larger population of P2rx7^-/-^ OT-II cells. Thus, *in vivo* studies indicate that P2X7R expression on T cells does not promote Th17 cell development but does suppress Th1 differentiation. Moreover, P2X7R expression can limit the overall T cell population through cell death, likely as a self-limiting mechanism to constrain inflammation.

**Figure 5.**
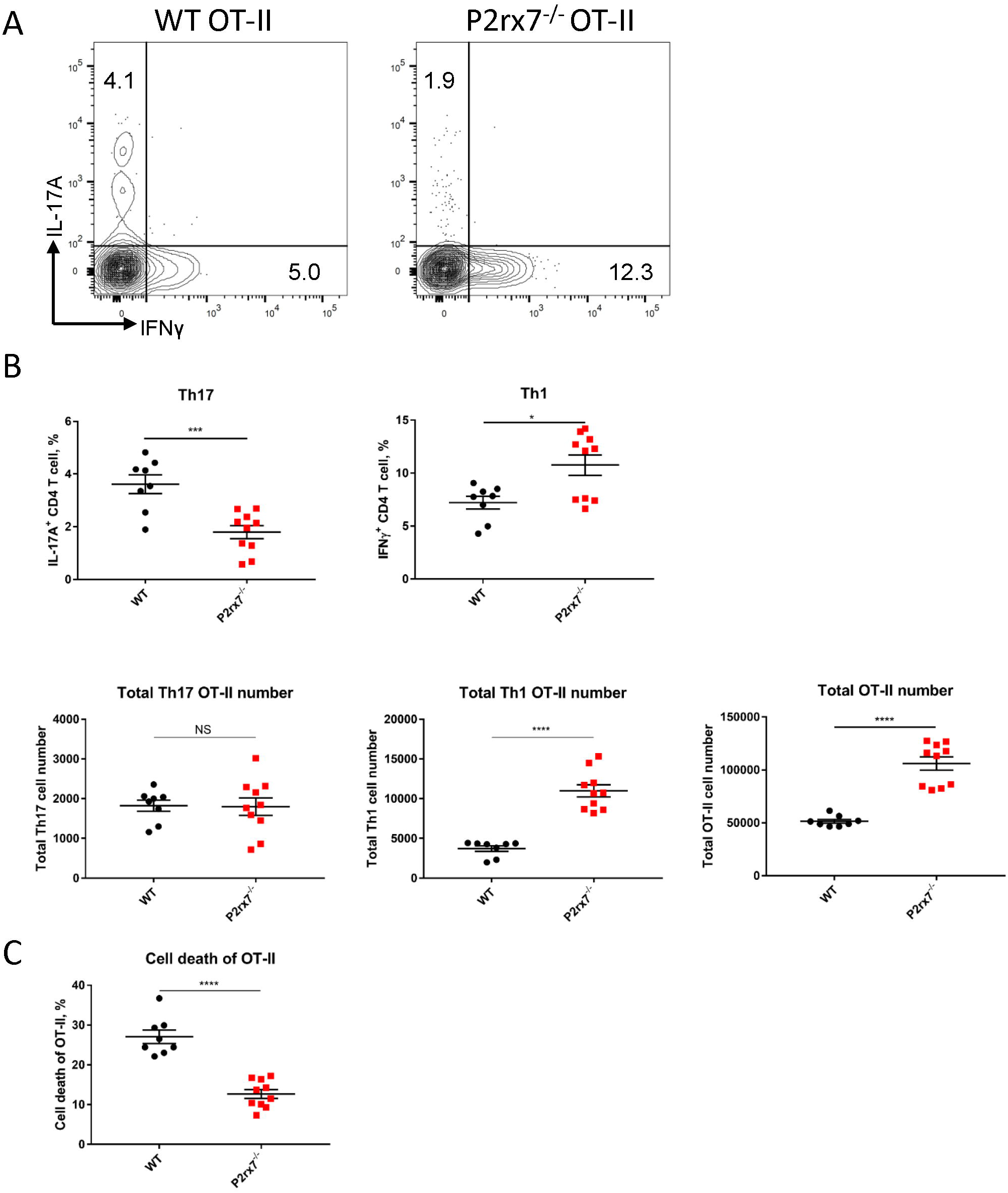
*In vivo* P2X7R expression on CD4^+^ T cells is not essential for Th17 differentiation. **(A-C)** Naive WT or P2rx7^-/-^ OT-II CD4^+^ T cells (both CD45.2) were adoptively transferred intravenously into congenic WT CD45.1 mice. The following day mice were immunized with OVA_323-339_/CFA. Inguinal LNs were collected and restimulated 7 days post-immunization. **(A)** Representative flow cytometric contour plots. **(B)** The percentage and total cell number of IL-17A^+^ and IFNγ^+^ OT-II T cells in the CD45.2 gate, were quantitated. **(C)** Cell death quantified by viability staining dye. Results shown as mean ± SEM. One representative of three independent experiments, n=19-20 total. *p<0.05, ***p<0.001, ****p<0.0001; NS, not significant.

## Discussion

Perhaps the earliest connection of ATP to IL-1β proteolytic maturation and release came from a study demonstrating that treatment of LPS-stimulated mouse peritoneal macrophages, with ATP as an apoptosis-inducing reagent, can induce the secretion of the mature form of IL-1β (24. Later studies demonstrated that the K+ efflux induced by ATP was critical for IL-1β processing in macrophages {Perregaux, 1994 #1220). ATP’s effect was also linked to the activation of the inflammasome and caspase-1, which cleaves pro-IL-1β to produce its active form (1). It is proposed that ATP provides the second signal with the first signal being the stimulation of pro-IL-1β production through NF-κB, provided by microbial components (25). P2X7R, as an ATP receptor, was introduced into this “Two signal model” almost at the same period (26–28). Notably, these studies were performed in macrophage-type cells and generalized to other cell populations. Our results herein demonstrate that BzATP stimulation of BMDCs can also induce IL-1β secretion. Likewise, human skin migratory (smi)DCs stimulated through the P2X7R expressed IL-1β and IL-6, which were blocked following the addition of P2X7R antagonists (11). Moreover, following P2X7R stimulation on BMDCs, we demonstrate that DCs can significantly increase the IL-17A production and Th17 differentiation by co-cultured naive CD4^+^ T cells. These findings are in line with our previous studies utilizing human smiDCs, in which we determined that purinergic signaling provokes innate cutaneous inflammatory responses, DC17 differentiation, and Th17 responses, in part, through the activation of mir-21 (11). Of note, in the absence of exogenously added BzATP, P2rx7^-/-^ BMDCs are capable of inducing comparable levels of IL-17A from co-cultured T cells with that of WT BMDCs, albeit at significantly lower levels, indicating that BMDCs likely have compensatory mechanisms for biasing Th17 responses. This notion is supported by our *in vivo* data that indicates that a cDC-specific P2X7R deficiency has a comparable percentage of IL-17A^+^ T cells and significantly higher IFNγ^+^ cells compared with P2X7R-sufficient cDCs. Thus, it appears that P2X7R expression on DCs can enhance Th17 development but is not necessary to induce Th17 cell development. These results do not rule out a role for radio-resistant LCs and other non-cDCs expressing P2X7R. In this regard, our previous studies have demonstrated that human LCs can induce Th17 differentiation (29). Though, systemic immunization is not likely affected by cutaneous LCs. Moreover, in most adaptive immune responses, including in our model, cDCs, especially CD11b^+^ cDCs, are thought to be responsible for Th17 CD4^+^ T cell responses, due to their higher expression of genes involved in MHC-II presentation compared to other cDC subtypes and are located at most environmental interfaces (30).

In the collagen-induced arthritis model, P2X7R is required for Th17 development as indicated following treatment with P2X7R antagonists (10). In contrast, there are studies using P2rx7^-/-^ mice that show P2X7R deficiency does not alleviate inflammation but enhances it. The models used in these studies are *Citrobacter rodentium* (Th17, Th1) or *Toxoplasma gondii* (Th1) infection, and the experimental autoimmune encephalomyelitis (EAE) model (Th17) (14–16). It is possible that the discrepancy is caused by the utilization of different models with different dependencies on Th17 responses. Another possible explanation is that P2X7R functions differentially on different tissues and cell types and whole-body knockout of this receptor will counteract its effects and mask the phenotype. Moreover, ATP and BzATP can both be metabolized to adenosine which is an anti-inflammatory agent that may also affect cell types differentially. Herein, our results demonstrate that global P2rx7^-/-^ mice can mount comparable Th17 and Th1 immune responses with that of WT mice. Thus, to further test the function of P2X7R on T cells we utilized *in vitro* co-cultures and P2rx7^-/-^ OT-II T cells. We demonstrate that P2X7R expressed by CD4^+^ T cells promotes Th17 development *in vitro* but P2X7R expression on CD4^+^ T cells is not essential *in vivo* for Th17 differentiation. However, we do demonstrate that Th1 responses are increased in the absence of P2X7R on CD4^+^ T cells. Importantly, in accordance with previous reports (3,22), we determined that P2X7R can induce T cell death *in vivo* resulting in the comparable total number of Th17 cells in WT and P2rx7^-/-^ OT-II cells. Thus, we hypothesize that Th17 cells are more sensitive to P2X7R induced cell cytotoxicity. Overall, our study indicates that P2X7R can enhance but is not required for Th17 differentiation. These findings contribute to our understanding of P2X7R and suggest celltype specific targeting strategies could improve the clinical efficacy of P2X7R therapeutics.

## Materials and Methods

### Mice

B6.129P2-P2rx7^tm1Gab^/J (P2rx7^-/-^), C57BL/6J, B6.SJL (CD45.1), BALB/cJ, OT-II, and zDC-DTR mice were purchased from The Jackson Laboratory. P2rx7^-/-^ OT-II mice were generated by crossing P2rx7^-/-^ with OT-II mice. CD45.1 OT-II mice were obtained by breeding B6.SJL with OT-II mice. Mice were maintained in specific pathogen-free conditions and all experiments were performed in accordance with Institutional Animal Care and Use Committee and the NIH guide for care and use of laboratory animals. Experiments and protocols were approved by the University of Pittsburgh’s IACUC.

### Adoptive transfer and immunization

Naive CD45.1^+^ OT-II CD4^+^ T cells were isolated with MACS (Mouse naive CD4^+^ T cell isolation kit from Miltenyi Biotec, Bergisch Gladbach, Germany) and transferred intravenously into congenic mice (10^5^ cell/mouse). The following day recipient mice were immunized subcutaneously at tail base with 20 μg OVA_323-339_ peptide (Anaspec, Fremont, CA) emulsified in CFA (BD biosciences, San Jose, CA) (or 20 μg KLH in CFA for some experiments). Seven days later, inguinal lymph nodes were collected and processed into single cell suspensions. For detection of cytokines in supernatants, cells were restimulated with cognate antigens for three days and then supernatants were collected for IL-17A and IFNγ detection with ELISA (Biolegend, San Diego, CA). For determination of IL-17A^+^ and IFNγ^+^ CD45.1^+^ OT-II CD4^+^ T cell percentages, cells were restimulated with 50 ng/ml PMA and 500 ng/ml ionomycin in the presence of GolgiPlug for 4 hours and stained for flow cytometry analysis.

### Antibodies and flow cytometry

The following antibodies were from BD: anti-CD16/CD32(2.4G2), BUV395 anti-CD3e(145-2C11), BUV737 anti-CD45.1(A20), PE anti-IFNγ(XMG1.2), Alexa Fluor 647 anti-IL-17A(TC11-18H10). The following antibodies were from Biolegend: Alexa Fluor 488 or PerCP/Cy5.5 anti-CD4(GK1.5), Brilliant Violet 605 anti- and CD45.2(104). Viability dye eFluor 450 was from eBioscience, San Diego, CA. For intracellular staining, surface stained cells were fixed and permeabilized with BD Cytofix/Cytoperm kit following manufacturer’s instructions and stained intracellularly with indicated cytokine antibodies. For indicated experiments, cell death was quantified by viability dye staining. For determination of absolute cell count cells were collected from each sample and mixed with 100 μl of AccuCheck Counting Beads (Life technologies, Carlsbad, CA). Absolute cell count was calculated for each sample based on the percentage and number of beads and sample volume as per manufacturer’s instructions. Cells were analyzed using an LSR II flow cytometer or the LSR Fortessa and FlowJo software was utilized for analysis (BD Biosciences).

### Bone marrow-derived DCs

Mouse bone marrow was isolated from femur and tibia. RBCs were lysed and T and B cells were depleted with antibodies (BD) plus rabbit complement (Cedarlane, Burlington, Canada). Cells were cultured at 1×10^6^ cells/ml in complete RPMI-1640 supplemented with 20 ng/ml GM-CSF and 5 ng/ml IL-4 (PeproTech, Cranbury, NJ) for 7 days (Replaced 70% of medium on Day 2 and Day 5). Loosely adherent BMDCs were collected and further purified with CD11C^+^ magnetic beads (Miltenyi Biotec, Bergisch Gladbach, Germany) to obtain a purity of >90%. BMDCs were stimulated with 100, 300, 500 μM of BzATP (SigmaAldrich, St Louis, MO) for 3h, 6h, or 24h as indicated, and supernatants were harvested for IL-6 and IL-1β ELISA (Biolegend, San Diego, CA). BMDCs were also utilized in the MLR and *in vitro* T cell differentiation assays.

### *In vitro* T cell differentiation

Naive CD4^+^ T cells from spleens of wild-type C57BL/6J and P2rx7^-/-^ mice were sorted with MACS columns (Miltenyi Biotec, Bergisch Gladbach, Germany). For cytokine-skewed Th17 differentiation, naive CD4^+^ T cells were stimulated with plate-bound 10 μg/ml anti-CD3 (Bio X Cell) and 2 μg/ml anti-CD28 (BD) in the presence of 20 ng/ml IL-6 (PeproTech), 5 ng/ml TGF-β1 (Biolegend), 20 ng/ml IL-23 (Miltenyi), and 20ng/ml IL-1β (PeproTech)(17). For Th1 differentiation, 20 ng/ml IL-12 (PeproTech) was used along with anti-CD3 and anti-CD28. After 4 days, cells were washed and restimulated with 50 ng/ml PMA and 500 ng/ml ionomycin in the presence of GolgiPlug for 4 hours and intracellularly stained for flow cytometric analysis. For experiments to track T cell proliferation, before culture in the skewing medium, naive CD4^+^ T cells were labelled with 5 μM CFSE (Molecular Probes, Eugene, OR) in PBS containing 5% FBS for 5 min at RT and washed 3 times.

T cell cocultures with BMDCs were performed as previously described (21), with modifications. Briefly, 2×10^4^ naive CD4^+^ T cells were sorted from wild-type C57BL/6J or P2rx7^-/-^ mice and 1×10^4^ BMDCs derived from wild-type C57BL/6J were cocultured in complete RPMI-1640 supplemented with 0.2 μg/ml anti-CD3, 100 ng/ml LPS (TLRgrade, from Enzo Life Sciences, Farmingdale, NY), and 1 ng/ml TGF-β1 (Biolegend) in roundbottom 96-well plates. Four days following coculture, cells were restimulated for 4 hours as described above and stained for flow cytometric analysis of IL-17A and IFNγ-producing CD4^+^ T cell.

### Mixed leukocyte reaction (MLR)

1×10^4^ BMDCs derived from wild-type C57BL/6J or P2rx7^-/-^ mice were cocultured with 1×10^5^ naive CD4^+^ T cells sorted from spleen of BALB/cJ mice in 200 ul of complete RPMI-1640 in round-bottom 96-well plates pre-coated with 5 μg/ml anti-CD3 for 4 days. Supernatants were harvested and IL-17A was measured with ELISA kit (Biolegend). For some experiments, BMDCs were pre-stimulated with 300 μM BzATP for 5 hours and washed before coculture.

### Mixed bone marrow chimera studies

Recipient C57BL/6J mice were irradiated with two doses of 550 rad, 3 hours apart. Mice were fed with Uniprim diet (Envigo, Indianapolis, IN) for two weeks. The following day, bone marrow was isolated from donor mice and RBCs and T cells were depleted. Cell density was similarly adjusted for bone marrow from each strain (between 2.0 ×10^7^ to 4.0×10^7^ cells/ml). Bone marrow cells from the two mouse strains were mixed 1:1 and 4 ×10^6^ to 8 ×10^6^ cells were injected intravenously per recipient mouse. Between 8 to 12 weeks after reconstitution, naïve CD45.1^+^ OT-II CD4^+^ T cells were injected into each chimera. The day following T cell transfer, chimeras were injected i.p. with either PBS or 1.25 μg Diphtheria Toxin (Sigma). To maintain DT induced DTR^+^ cell depletion, DT was injected every other day for the duration of experiments. Chimeras were immunized with 100 μg OVA_323-339_ peptide in CFA at tail base 24 hours after the first DT injection. 7 days after immunization, inguinal LNs were collected and made into a single cell suspension, which was restimulated for 4 hours as described above and stained for flow cytometry analysis of IL-17A and IFNγ-producing CD45.1^+^ CD4^+^ T cell. Counting beads were used to determine absolute cell count.

### Statistics

All results were analyzed with GraphPad Prism7 Software (San Diego, CA). Statistical differences were obtained by using Student’s t test for two groups and two-way ANOVA for multiple groups. *p<0.05, **p<0.01, ***p<0.001, ****p<0.0001.

## Acknowledgements

Research reported in this publication was supported by the National Institute of Arthritis and Musculoskeletal and Skin Diseases of the NIH under Award Number R01AR067746 (to A.R.M.). The content of this manuscript is solely the responsibility of the authors and does not necessarily represent the official views of the National Institutes of Health. The authors declare no conflict of interest.

