## Supplemental Figure 1 for "P2X7 receptor signaling is not required but can enhance Th17 differentiation"

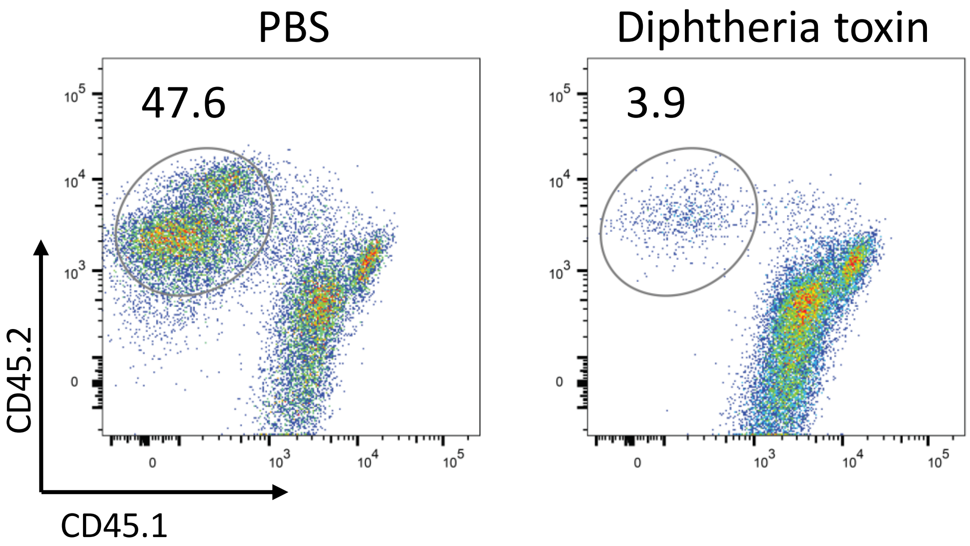


**Supplementary Figure 1.** Successful reconstitution of mixed bone marrow and DT depletion of classical DCs only from zDC-DTR bone marrow (CD45.2) was shown for the mixed bone marrow chimeras experiment in Fig3.B-C. Percentage of cDC (gated on $Lin^-/CD11C^+/MHC\ II^{high}$) (Meredith et al., 2012) derived from CD45.2 zDC-DTR bone marrow cells is shown.
